# Matrix stiffness-induced IKBKE and MAPK8 signaling drives a phenotypic switch from DCIS to invasive breast cancer

**DOI:** 10.1101/2025.04.16.649065

**Authors:** Feifei Yan, Sara Göransson, Helene Olofsson, Christos Vogiatzakis, Anagha Acharekar, Staffan Strömblad

**Affiliations:** Department of Medicine Huddinge, Karolinska Institutet, SE-141 83 Huddinge, Sweden; Clinical Trials Office, Centrum för Kliniska Cancerstudier, Tema Cancer, Karolinska Universitetssjukhuset, SE-171 76 Stockholm, Sweden

## Abstract

Ductal carcinoma in situ (DCIS) is not life threatening unless it transitions into invasive breast cancer (IBC). However, although breast cancer cell exposure to matrix stiffening in vitro phenotypically mimics the DCIS to IBC switch, the molecular changes driving this switch remains unclear. Here, breast cancer cell kinome activity profiling suggested matrix stiffness-upregulation of 53 kinases, among which 16 kinases were also regulated by integrin β1. Functional validation identified matrix stiffness-activation of inhibitor of nuclear factor kappa-B kinase subunit epsilon (IKBKE) and mitogen-activated protein kinase 8 (MAPK8) signaling as critical for the stiffness-driven IBC phenotype, including for cell proliferation. The IKBKE-inhibitor Amlexanox, clinically utilized for aphthous ulcers, as well as the MAPK8 inhibitor JNK-IN-8, reinstalled the DCIS-like phenotype of breast cancer cells on high matrix stiffness. This suggests that IKBKE and/or MAPK8 inhibitors could enhance the arsenal of treatments to prevent or treat breast cancer.

## Introduction

Breast cancer is the most commonly diagnosed cancer and the leading cause of cancer deaths in women[1]. It is a heterogeneous disease with many subtypes showing immensely different disease manifestations and clinical outcomes[2]. Screening mammography was introduced in many parts of the world following randomized controlled studies reporting around a 30% reduction in risk of dying from breast cancer in the group of women invited to screening[3, 4]. This widespread adoption of screening mammography has led to a marked increase in detected cases of preinvasive lesions, including ductal carcinoma *in situ* (DCIS)[5]. At this early stage of the disease, neoplastic cells are still confined to the ductolobular system of the breast and DCIS is often regarded as an early form of breast cancer even though most of these preinvasive lesions never progress to IBC[6]. However, the molecular mechanisms that drive neoplastic cells to invade the surrounding stroma remain unclear.

Several studies indicate that DCIS and invasive cancers display very similar molecular profiles[7-11], suggesting that copy number alterations and gene expression changes occur already at the DCIS stage. In contrast to this, larger differences have been found in the respective microenvironments[8, 12, 13]. Moreover, changes in collagen organization at the tumor border facilitates local cancer cell invasion[14] and can be used to predict breast cancer survival[15, 16]. The progression from normal tissue to DCIS and further to invasive ductal carcinoma (IDC) involves increased deposition and organization of collagen leading to progressively stiffer extracellular matrix (ECM)[17]. Stiffening of the ECM, a hallmark of many solid cancers, disrupts the tensional balance[18] and fuels cancer cell proliferation, invasion, and drug resistance and hence cancer progression[19, 20]. This is because components of the ECM serve as ligands for cell surface receptors, like integrins, that sense and transduce mechanical properties of the environment to induce a functional cellular response[21], often via changes in kinase signaling. Classical mechanosensitive signaling downstream of integrins, e.g., FAK/Src, RhoA/ROCK, MAPK, and PI3K, are pro-survival and pro-proliferative and contribute to disease progression[22-24]. Consequently, targeting this matrix stiffness-mediated signaling may be a strategy to prevent breast cancer invasion and metastasis[25].

We previously showed a correlation between matrix stiffness-induced mRNA changes and gene sets describing the difference between DCIS and co-occurring IDC in breast cancer patients[26]. This suggests matrix stiffness as a facilitator in the transition from DCIS to IDC and calls for further studies to elucidate signaling that governs the stiffness-induced malignant phenotype. However, given that the transition from DCIS to invasive cancer displays limited alterations of mRNA expression, it is important to define other levels of molecular regulation during this transition.

Here, we combined the *in vitro* hydrogel-based culture system used in the transcriptome study with a peptide array to profile changes in kinase activity upon the stiffness-driven transition from a DCIS to an IDC phenotype. We validated the function of the predicted kinases using RNAi-based screening and identified the IKBKE and MAPK8 kinases as mechanoregulated proteins being potential targets for development of novel breast cancer treatment.

## Results

### Reversal of matrix stiffness-induced phenotypic switch in MCF10CA1a cancer cells by integrin β1 inhibition

MCF10CA1a (CA1a) is a fully malignant human breast cancer cell line generated from the non-malignant MCF10A cell line[27]. CA1a cells form undifferentiated tumors when implanted subcutaneously and lung metastasis when injected into the tail vein of immunocompromised mice[27]. To mimic a normal mammary gland microenvironment, we cultured these cells on compliant hydrogels conjugated with reconstituted basement membrane (rBM). On hydrogels with stiffness mimicking normal breast tissue (0.4-0.5 kPa) these cells formed mainly non-invasive DCIS-like structures that contrasted dramatically to the invasive growth pattern observed on stiffness similar to that of breast cancer(5-8 kPa) (Fig. 1a). The cellular phenotype on low stiffness exhibited an intact basement membrane, as indicated by laminin staining (Fig 1b), with properly polarized cells as marked by integrin β4 localization to the outer rim (Fig. 1b). These results are in agreement with early studies showing that cues from a normal microenvironment can revert malignant cells to a more normal phenotype[26, 28] as well as seminal work describing the effects of matrix stiffness on breast cancer progression[18, 29].

**Fig. 1:**
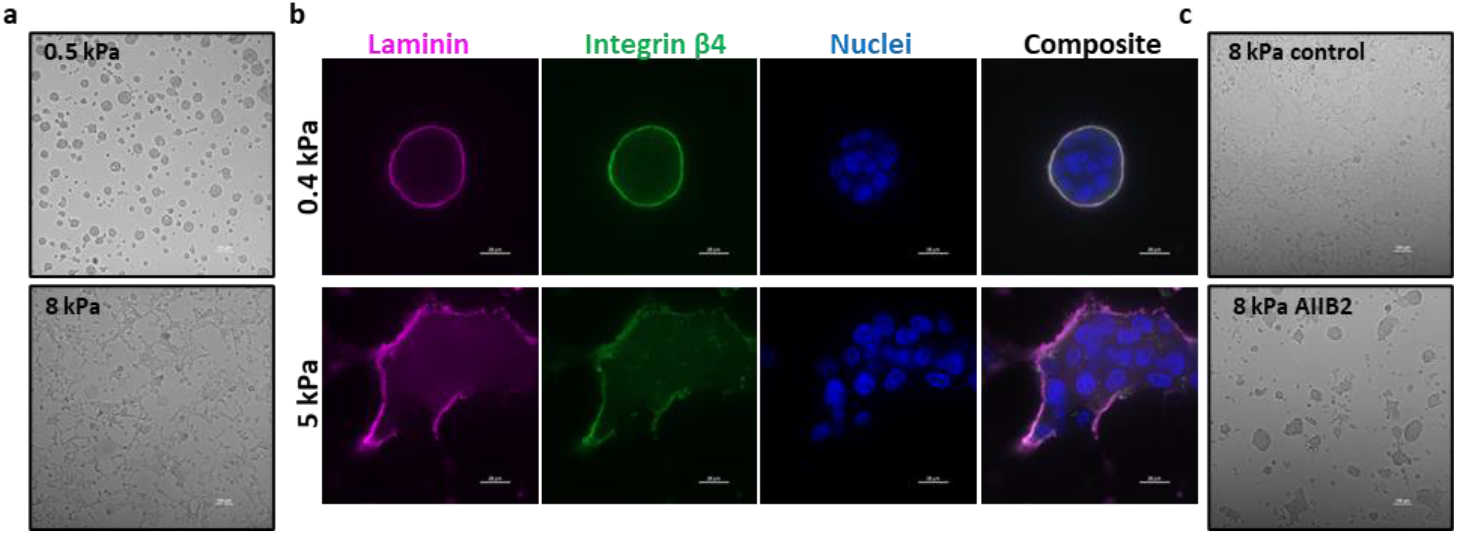
Matrix stiffness induced switch in MCF10CA1a (CA1a) cancer cell phenotype reversed by integrin β1 inhibition. **a**, Representative phenotype of CA1a cells on rBM-conjugated hydrogels of different stiffness mimicking normal breast tissue stiffness (0.5 kPa) or breast cancer stiffness (8 kPa). Scale bar = 100 µm. **b**, Representative spinning disc confocal immunofluorescence images of laminin (pink), integrin β4 (green) and nuclei (blue) in CA1a cells on low (top row) and high (bottom row) stiffness hydrogels. Scale bar = 20 µm. **c**, CA1a cells on high stiffness hydrogels treated with anti-integrin β1 monoclonal AIIB2 antibody or vehicle alone. Scale bar = 100µm.

Increased integrin signaling, as a result of ECM stiffening, is known to promote tumor-like behavior in mammary epithelial cells[18], and blocking integrin beta 1 signaling normalizes this malignant behavior in culture[18] as well as *in vivo*[30]. To evaluate the role of integrin beta 1 in our culture system we used the anti-integrin β1 monoclonal antibody AIIB2 to specifically block this signaling. As expected, treating CA1a cells on high stiffness with the AIIB2 antibody for 18-20 h induced a reversion of the malignant phenotype (Fig. 1c).

### Profiling of the mechanosensitive kinome by peptide chip arrays

To elucidate the signaling governing the stiffness-induced transition to an invasive phenotype, we performed tyrosine kinase (PTK) and serine/threonine kinase (STK) activity profiling using commercially available peptide chip arrays from PamGene[31] (Fig 2a). These arrays contain 196 tyrosine and 144 serine/threonine bait peptides, respectively, and the pattern of phosphorylation following incubation with cell lysate is used to predict the activity of specific kinases. Lysates were prepared in biological triplicates from CA1a cells cultured on soft (0.5 kPa) or stiff (8 kPa) rBM-conjugated hydrogels for 3 d. To specifically elucidate stiffness signaling transduced via integrin beta 1, anti-integrin β1 mab AIIB2 was added for the last 20 h of culture to one group of cells on stiff gels (Fig. 2a). While all experimental groups contained three biological replicates, unfortunately, one array on STK chip #2 broke during the run, leaving us with only two AIIB2 replicates for STK. Principle component analysis (PCA) based on all peptides passing quality control (QC) in the respective array (PTK and STK) successfully separated low and high stiffness samples in the first component (Fig. 2 b). Out of the 79 QC PTK peptides, 59 were significantly more phosphorylated on high stiffness (p-value < 0.05) (Fig. 2c, Supplementary data 1). On the STK array, 67 out of 108 QC peptides were differentially phosphorylated (p-value < 0.05) (Fig. 2d, Supplementary data 2). To evaluate if any stiffness-induced phosphorylation occurred downstream of integrin β1, we performed PCA analysis of high stiffness samples and AIIB2 treated samples using the peptides that were more significantly regulated (p < 0.01) by stiffness. This revealed a separation of the two groups in the second component, suggesting that the phosphorylation pattern generated by increased stiffness is changed when inhibiting integrin β1 signaling (Fig. 2e). Volcano plots showing the log2 fold change and p-value distributions of the AIIB2 versus high stiffness comparison show that the serine/threonine phosphorylation was more affected by AIIB2 treatment than tyrosine phosphorylation (Fig. 2f, g). Four PTK peptides and eleven STK peptides were differentially phosphorylated upon AIIB incubation with a p-value below 0.05 (Supplementary data 3). Differentially phosphorylated peptides were subsequently used to predict kinase activity by computational modelling of differentially phosphorylated peptides, applying a specificity threshold (mean specificity score > 1). Among the kinases predicted to be activated by stiffness are well-characterized mechanosensitive signaling components such as ERK1 and Src [32] and also kinases not previously linked to mechanosignaling, including IKBKE and ZAP70 (Fig. 2h). Not surprisingly, inhibition of integrin beta 1 attenuated ERK, JNK, and p38 serine/threonine kinases, known integrin signaling mediators (Fig. 2i)[32, 33]. The prediction of a minor upregulation of tyrosine kinase activity upon AIIB2 treatment is somewhat surprising but may represent a feedback mechanism or indicate that other integrin isoforms and/or growth factor receptors are important for tyrosine kinase activation in our model (Fig. 2i).

**Fig. 2:**
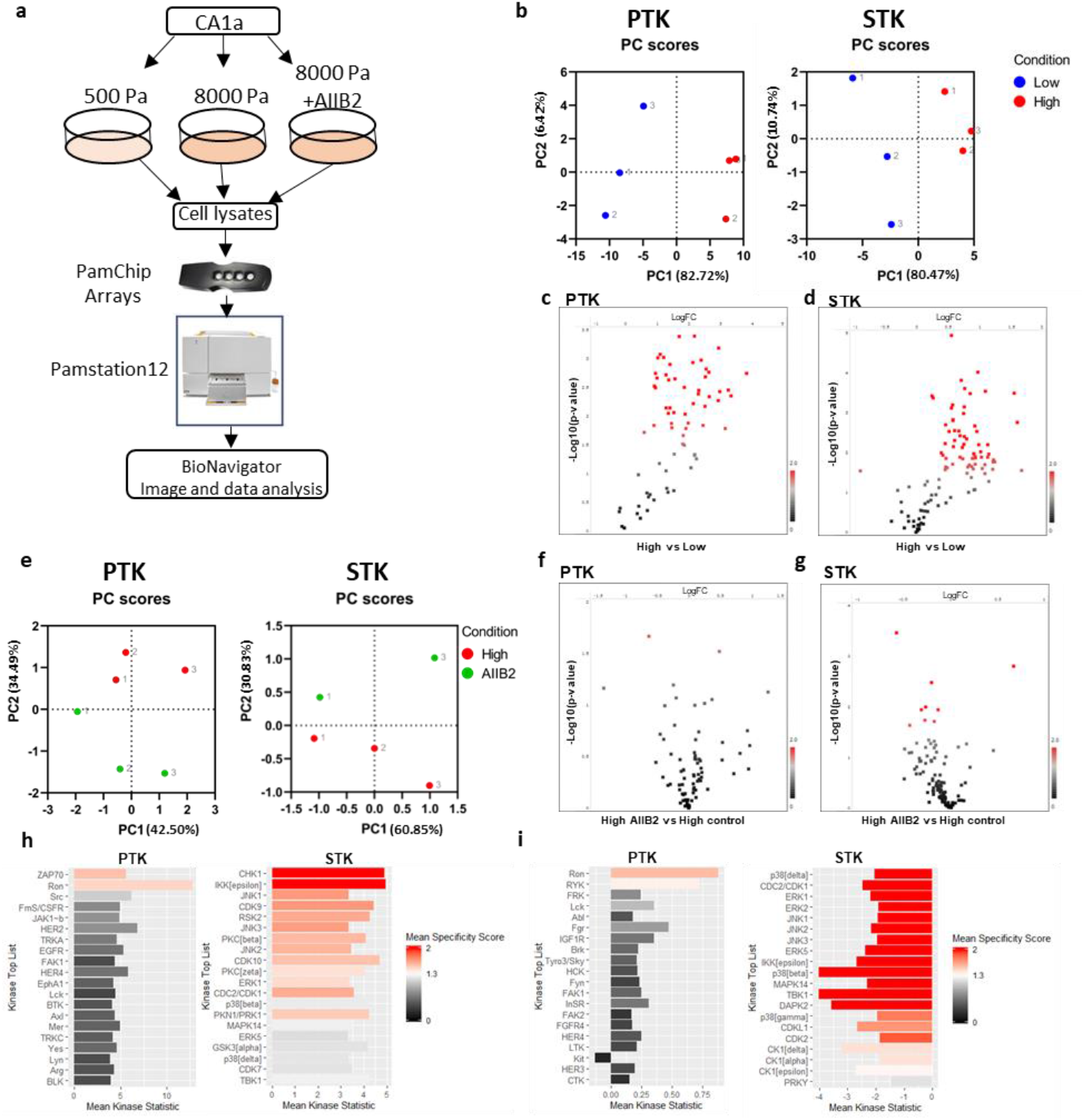
Kinome activity profiling reveals matrix stiffness-induced changes in tyrosine and serine/threonine peptide phosphorylation partially dependent on integrin β1. **a**, Experimental setup for peptide array-based kinome profiling of CA1a cells on hydrogels. **b**, Principal component analysis (PCA) of low and high stiffness samples using all QC peptide phosphorylation data from PTK (left) and STK (right) chip arrays. Colour indicate condition and number indicate replicate **c, d**, Volcano plots of peptide phosphorylation data from high stiffness vs low stiffness PTK (c) and STK (d) chip arrays assessed by unpaired t-test. Red spots are peptides with p < 0.05. **e**, PCA analysis of high stiffness and high+AIIB2 treated samples using only the most differentially phosphorylated peptides (p < 0.01) from the low versus high stiffness comparison. **f, g**, Volcano plots of peptide phosphorylation data from high stiffness AIIB2 treated vs high stiffness control treated PTK (f) and STK (g) chip arrays assessed by unpaired t-test. Red spots are peptides with p < 0.05. **h, i**, Kinase score plots for high stiffness versus low stiffness (h) or for AIIB2 versus control (i) from PTK (left) and STK (right) peptide chip array data. Upstream kinases are predicted from differentially phosphorylated peptides using the proprietary Bionavigator software, with a mean specificity score >1.

### Peptide array-based kinome activity profiling predicts kinases regulated in the stiffness-induced switch to the malignant breast cancer phenotype

In addition to the kinase regulation predicted by modelling described above (Fig 2h-i), we then added also peptides in the array that were derived from known kinases and exhibited statistically discernible altered phosphorylation. This way, we identified a total of 53 kinases predicted to be upregulated by increased matrix stiffness (Fig 3a, Supplementary data 4) (p < 0.05). A comparison of the kinases whose activity was predicted to be upregulated by stiffness and kinases inhibited by AIIB2 treatment revealed an overlap of 16 kinases (Fig 3a, Supplementary data 4). To evaluate the possible function of these 16 kinases in driving the malignant phenotype on high stiffness, we performed a limited RNAi screen (Fig. 3b). Kinases were knocked-down using siRNA pools specific for each kinase and transfected cells were plated on high stiffness rBM-conjugated hydrogels for 4 d before assessing the phenotype using widefield imaging of the entire hydrogel surface followed by image analysis (Fig. 3b). Negative control (MOCK and scrambled siRNA) and positive control (FAK siRNA) were included in all hydrogel plates. Knockdown of FAK was used as a positive control due to its established role as a mediator of stiffness-induced phenotype[34, 35]. A kinase was considered as a mediator of the matrix stiffness-induced malignant behavior when its knockdown induced a change in the stiffness phenotype similar to FAK knockdown (Fig. 3c). The area fraction of cell clusters was calculated as the ratio between the total area and the area of the segmented objects and normalized to the area fraction of MOCK in each plate. This analysis indicated that the knockdown of 2 out of the 16 kinases induced a reversion of phenotype similar to FAK knockdown, including IKBKE and MAPK8 (Fig. 3c, d).

**Fig. 3:**
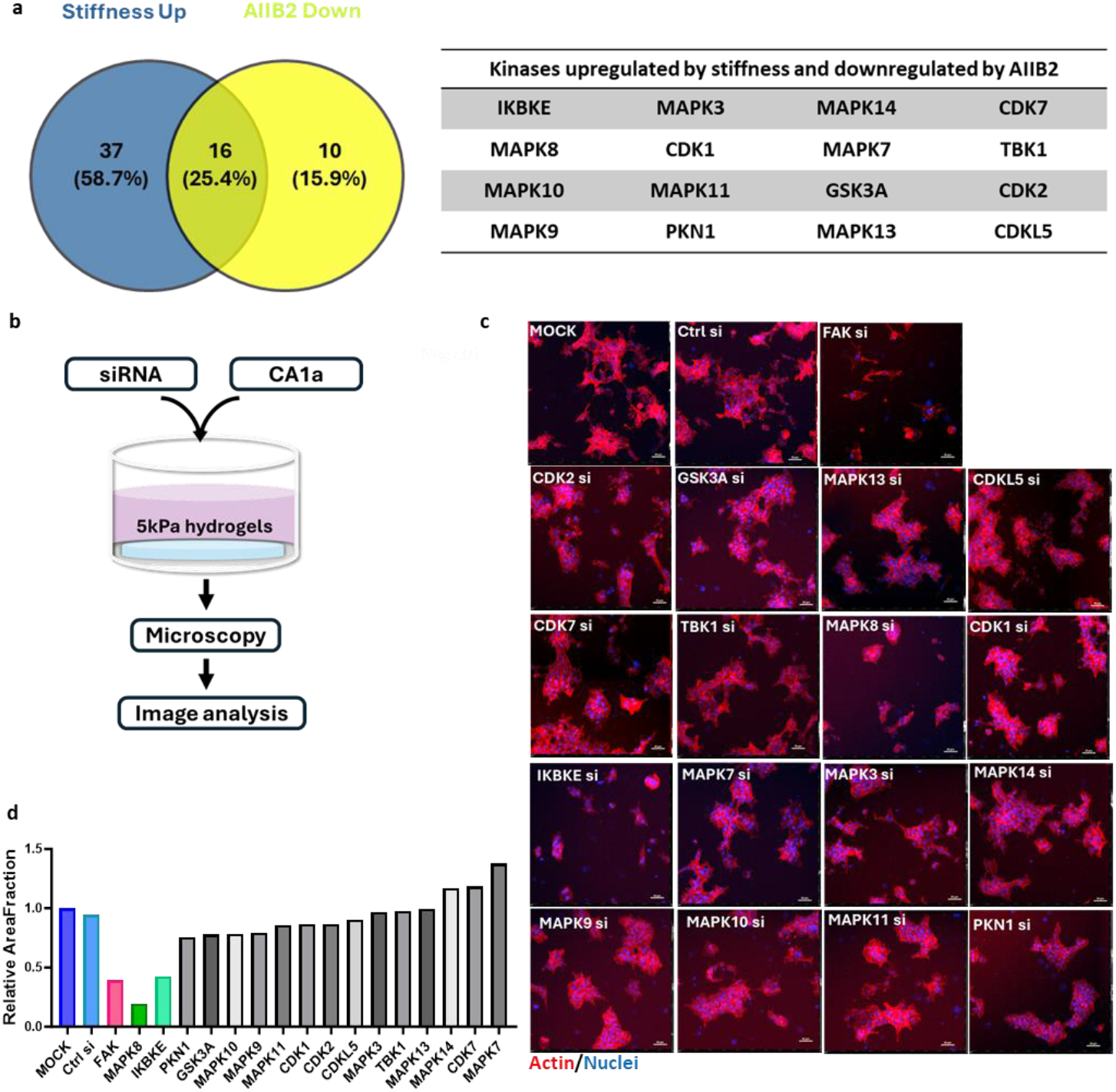
MAPK8 and IKBKE mediate the stiffness-induced breast cancer phenotype. **a**, Venn diagram (left) and table (right) showing overlap between kinases predicted to be upregulated by stiffness and downregulated by AIIB2 treatment. b, Workflow for the siRNA-based screen used to test the function of predicted kinases, and all conditions were run in duplicates. **c**, Representative phenotype of CA1a cells on high stiffness hydrogels transfected with siRNA as specified. FAK siRNA serves as positive control. Images are single tiles from 6×6 montages. Scale bar = 50 µm. **d**, Area fraction of segmented cell clusters expressed relative to MOCK (Gene of interest/MOCK). The area fraction is calculated as the ratio between the total area and the area of the thresholded objects. Positive control (FAK siRNA) is shown in red. Data represents the average of two technical replicates. Genes whose knockdown results in a similar or lower relative area fraction as the positive control are considered hits, including MAPK8 and IKBKE.

### IKBKE signaling drives a matrix stiffness-induced transition from DCIS to IDC phenotype

One of the identified kinases, inhibitor of nuclear factor kappa-B kinase subunit epsilon (IKBKE), a non-canonical I-kappa-B kinase, has been identified as a breast cancer oncogene amplified and overexpressed in more than 30% of breast carcinomas[36]. However, the potential relation of IKBKE to mechanosignaling has remained unclear. High matrix stiffness (5 kPa) promoted higher IKBKE protein expression in human breast cancer CA1a (Fig. 4a) and HCC1143 cells (Supplementary Fig. 1a), as compared to low stiffness (0.5 kPa). Transfection of CA1a cells with an IKBKE siRNA pool resulted in a phenotypic reversion from invasive breast cancer to DCIS and decreased cell proliferation (Fig. 3c, 3d and 4b). The knockdown efficiency of the four individual IKBKE siRNAs, which are mixed to form the IKBKE siRNA pool, was tested in human breast cancer HCC1143 cells (Supplementary Fig. 1d). HCC1143 cells transfected with each of two individual IKBKE siRNAs with confirmed knockdown efficiency showed a reversal of the stiffness IBC phenotype to the DCIS phenotype of low stiffness matrix (Fig. 4c, top row) as well as an inhibition of proliferation (Fig. 4c, bottom row). The role of IKBKE signaling for the stiffness-induced phenotype was further tested using a pharmacological inhibitor of IKBKE [37], Amlexanox, a dual IKBKE/TBK1 inhibitor that is clinically used for the treatment of recurrent aphthous ulcers and asthma[38, 39]. Treatment of CA1a cells on high stiffness with Amlexanox reverted the stiffness-induced phenotype (Fig. 4d, top row) and impaired cell proliferation (Fig. 4d, bottom row). Further validation of the IKBKE inhibitor was performed in HCC1143 cells, and also in these cells, the Amlexanox treatment inhibited the stiffness-induced malignant breast cancer cell phenotype (Fig. 4e, top row) and proliferation (Fig. 4e, bottom row). These findings indicate that mechano-regulated IKBKE signaling may promote breast cancer progression, and that targeting of IKBKE could be a potential strategy for prevention and/or treatment of breast cancer.

**Fig. 4:**
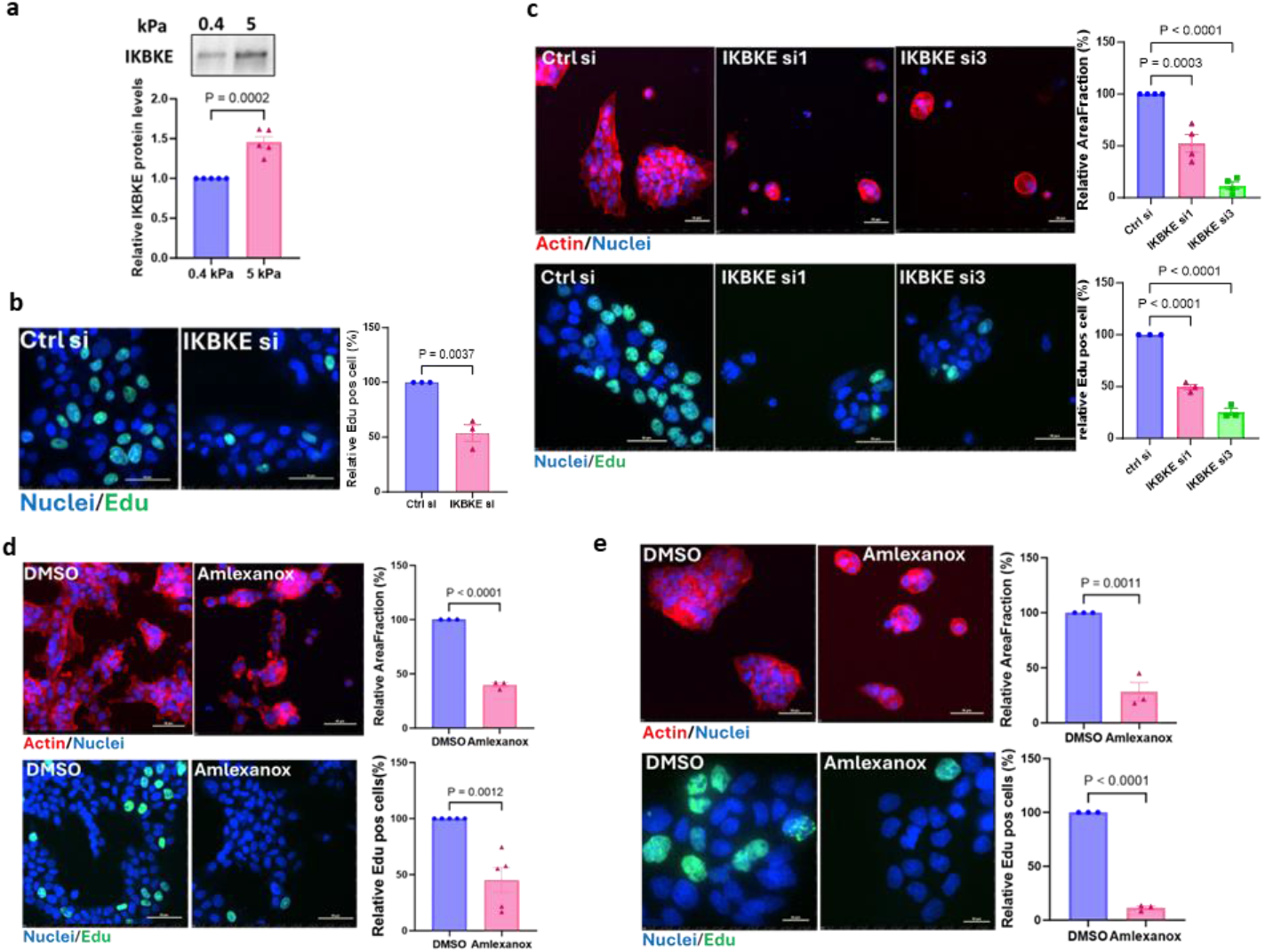
Mechano-regulated IKBKE signaling controls breast cancer cell growth. **a**, Immunoblot of IKBKE in CA1a cells (left) cultured on 0.4 kPa or 5 kPa hydrogels as indicated. Densitometric analysis shows IKBKE levels normalized to loading controls (ponceau S staining, Supplementary Fig. 2a) and expressed relative to the low stiffness level (right). Data are presented as mean ± S.E.M., with p-values according to an unpaired t-test. **b**, Representative immunofluorescent images of Edu staining in CA1a cells grown on 5 kPa transfected with IKBKE or control siRNA pool (left) and quantification of three biological repeats (right), each with more than one thousand cells. Scale bar = 50 µm. Data are mean ± S.E.M and p-values according to an unpaired t-test. **c**, Representative images of actin (top row) and Edu staining (bottom row) in HCC1143 cells treated with IKBKE or control siRNA (left). Scale bar = 50 µm. Area fraction of segmented cell clusters expressed relative to control siRNA. Quantification of three or more biological repeats (right). Data are mean ± S.E.M and p-values derived by one-way Anova followed by Dunnett’s multiple comparison test. **d, e**, Representative spinning disc confocal images of CA1a (d) and HCC1143 (e)cells on 5 kPa treated with 180 µM Amlexanox or vehicle for 48 h (left). Quantification of actin staining (top row) and Edu staining (bottom row) from three or more independent experiments (right). Scale bar = 50 µm. Relative area fraction as in (c). Data and statistics as in (b).

### Matrix stiffness-induced MAPK8 activity is a critical driver of breast cancer progression *in vitro*

The other identified kinase, MAPK8, also known as c-Jun N-terminal kinase 1 (JNK1), may play a dual and context-dependent role in cancer, with either tumor promoter or tumor suppressor properties depending on the cancer type and cellular environment[40-43]. In breast cancer, MAPK8 activation enhances the expression of ECM components such as osteopontin and tenascin C, which drive metastasis and therapy resistance by reinforcing the chemoresistant metastatic niche[44]. Additionally, MAPK8 stress signaling is hyper-activated in fibroblasts residing in high density breasts and tumor stroma where it contributes to fibrosis, inflammation, and the maintenance of cancer stem cell traits[45]. Despite its established role in ECM remodeling and tumor progression, the possible regulatory links from ECM stiffness to MAPK8 activity in breast cancer remain unclear. Interestingly, immunoblot analysis demonstrated a stiffness-dependent upregulation of MAPK8 activity without a concomitant increase in MAPK8 protein expression in CA1a cells (Fig. 5a) as well as in HCC1143 cells (Supplementary Fig. 1b and 1c). Knockdown of MAPK8 using an siRNA-pool significantly inhibited the invasive breast cancer phenotype (Fig. 3c and 3d) and cell proliferation (Fig. 5b) in CA1a human breast cancer cells. Additionally, two individual MAPK8 siRNAs with confirmed knockdown efficiency (Supplementary Fig. 1e) reversed the stiffness-induced phenotype (Fig. 5c, top row) and suppressed cell growth proliferation (Fig. 5c, bottom row) in HCC1143 human breast cancer cells. The role of MAPK8 activity was further tested using the selective MAPK8-inhibitor JNK-IN-8[46]. JNK-IN-8 effectively reverted the stiffness-induced phenotype and proliferation in both CA1a (Fig. 5d) and HCC1143 breast cancer cells (Fig. 5e). Collectively, these findings indicate that matrix stiffness-induced MAPK8 activity promote breast cancer progression and suggest that MAPK8 signaling represents a candidate for the development of new breast cancer treatment modalities.

**Fig. 5:**
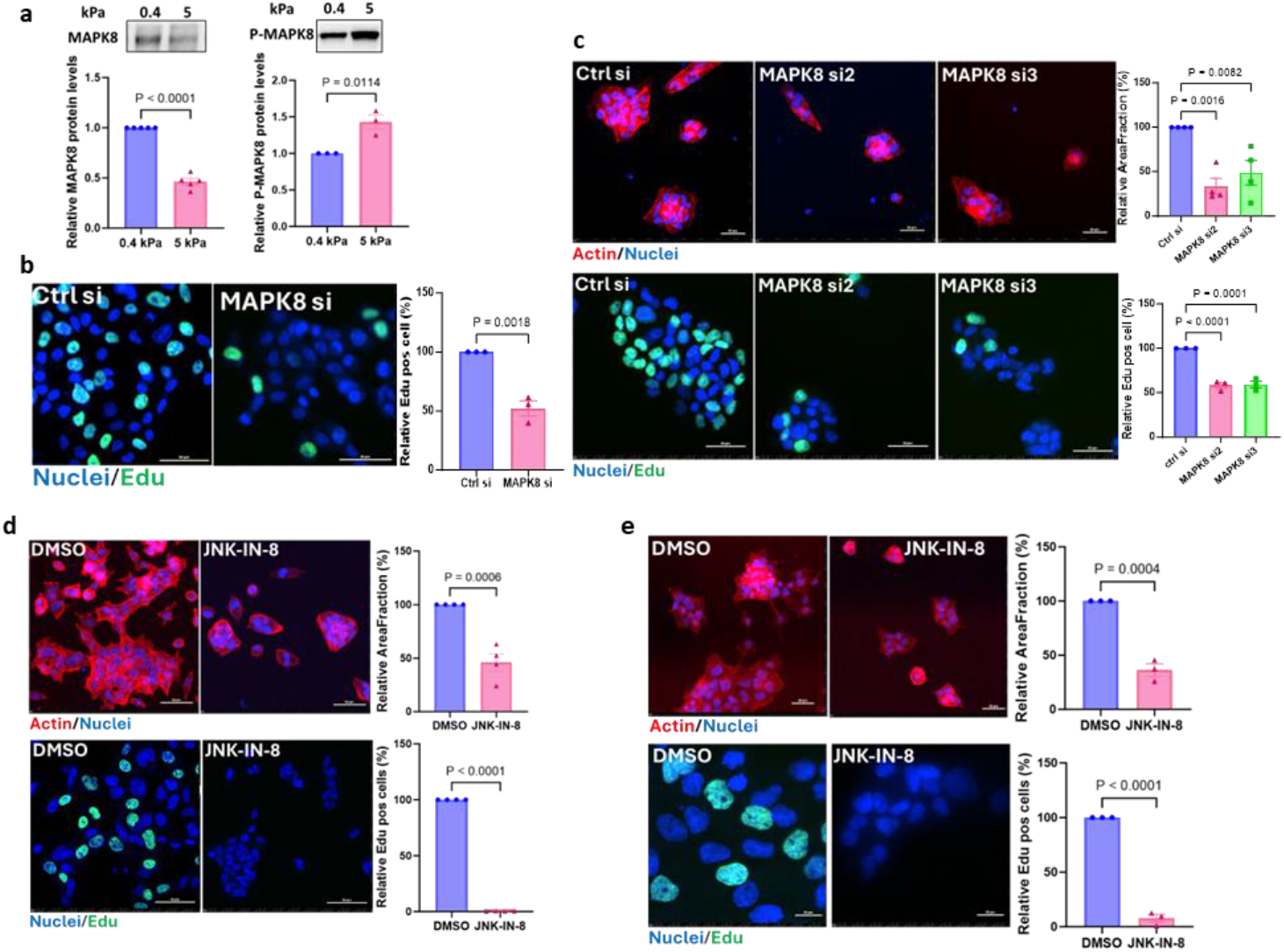
Matrix stiffness-induced MAPK8 activity is critical for breast cancer cell proliferation *in vitro*. **a**, Immunoblot analysis of MAPK8 (left), and phospho-MAPK8 (right) of CA1a cells cultured on 0.4 kPa or 5 kPa hydrogels. Densitometric analysis shows MAPK8 and phospho-MAPK8 levels normalized to loading controls (ponceau S staining, Supplementary Fig. 2a and 2c) and expressed relative to the low stiffness levels. Data are presented as mean ± S.E.M., with p-values according to an unpaired t-test. **b**, Representative Edu staining images of CA1a cells grown on 5 kPa transfected with MAPK8 or control siRNA pool (left) and quantification of three biological repeats (>1000 cells each; right). Scale bar = 50 µm. Data are mean ± S.E.M and p-values according to an unpaired t-test. **c**, Representative images of actin (top row) and Edu staining (bottom row) in HCC1143 cells treated with MAPK8 or control siRNAs (left). Quantification of area fraction of segmented cell clusters relative to control siRNA of three biological repeats (right). Scale bar = 50 µm. Data are mean ± S.E.M and p-values derived by one-way Anova with Dunnett’s multiple comparison test. **d, e**, Spinning disc confocal images of breast cancer cells on 5 kPa treated with 5 µM (CA1a, d) or 10 µM (HCC1143, e) JNK-IN-8 or vehicle for 24 h(left). Quantification of actin staining (top row) and Edu staining (bottom row) of three or more independent experiments(right). Scale bar = 50 µm. Relative area fraction as in (c). Data and statistics as in (b).

## Discussion

We used a commercially available peptide array in combination with siRNA-based functional validation and identified the kinases IKBKE and MAPK8 as mechano-regulated proteins critical for the matrix stiffness-induced phenotypic switch from DCIS to invasive breast cancer. An increased understanding of the molecular mechanisms driving the pre-invasive to invasive transition may be of great importance as it could generate biomarkers of progression as well as drug targets for invasive breast cancer.

Increased expression of IKBKE in cancer has been detected both with and without corresponding copy number gain[36, 47] and several recent studies implicate this kinase in breast cancer progression and metastasis[36, 47-49]. Similarly, our study points to IKBKE signaling in breast cancer progression, specifically in driving DCIS to IDC transition, and further suggests that IKBKE is activated as a result of augmented mechanical signaling, adding yet another layer of regulation of this kinase in breast cancer cells. The IKBKE/TBK1 inhibitor, Amlexanox, can repress cancer cell proliferation and invasion[50-53] and has shown efficacy in different experimental tumor models[54]. In line with these studies, our data suggests that Amlexanox is effective in reverting the malignant behavior induced by matrix stiffness. Future clinical trials may evaluate the repurposing of Amlexanox as well as the efficacy of other IKBKE inhibitors in breast cancer treatment. The anti-cancer effects of Amlexanox have been attributed to the inhibition of IKBKE and TBK1 but also to the inhibition of other proteins like FGF-1 and S100 proteins[54]. The effects of the dual kinase inhibitor Amlexanox in reverting the stiffness-induced phenotype resemble the effect we observe with IKBKE RNAi, but it remains to be investigated to what extent the effect of Amlexanox is specifically due to IKBKE inhibition.

MAPK8 signaling plays a multifunctional role in breast cancer, influencing tumor progression, metastasis, therapy resistance, and stem cell regulation[43-45]. MAPK8 activation remodels the tumor microenvironment, promoting metastasis and therapy resistance[44]. High breast density displays upregulated MAPK8 activity in fibroblasts and the tumor stroma[45]. We here provide direct evidence that increased matrix stiffness promotes MAPK8 activity without detectable changes in its protein expression. The MAPK8 inhibitor JNK-IN-8 has demonstrated therapeutic potential, since it synergizes with lapatinib to induce cell death in triple-negative breast cancer (TNBC) cells and significantly improves survival in mouse models with TNBC xenografts[55]. Additionally, JNK-IN-8 enhances the efficacy of FOLFOX chemotherapy in pancreatic ductal adenocarcinoma[46]. Another MAPK8 inhibitor, WBZ_4, has shown strong antitumor effects in ovarian cancer models both *in vitro* and *in vivo*[56]. Consistent with these findings, our data indicate that JNK-IN-8 significantly reduces breast cancer growth and reverses the stiffness-induced invasive breast cancer phenotype. JNK-IN-8 is highly potent against MAPK8 (JNK1) and also irreversibly inhibits JNK2 and JNK3. Further studies are therefore needed to elucidate if the effects of JNK-IN-8 in breast cancer cells are specifically due to MAPK8 inhibition. Clinical trials will be essential to evaluate the efficacy of JNK-IN-8, or other relevant MAPK8 inhibitors, as a potential therapeutic agent for breast cancer.

In addition to the validated IKBKE and MAPK8 kinases, we predicted other kinases previously known to work downstream of matrix stiffness and integrin signaling, providing validity to the peptide array as a tool for kinome profiling. In conclusion, we show that matrix stiffness activates several tyrosine and serine/threonine kinases and validated two of these kinases, IKBKE and MAPK8, as critical for a malignant breast cancer cell phenotype. This suggests that IKBKE and/or MAPK8 inhibitors could play a role in future breast cancer prevention and treatment.

## Methods

### Cell culture

MCF10CA1a cells were obtained from the Barbara Ann Karmanos Cancer Institute, Michigan, United States. Cells were maintained in DMEM/F12 (Gibco) media supplemented with 5% horse serum (Gibco), 10 mM HEPES (Gibco), and 0.029 M sodium bicarbonate (Gibco). HCC1143 cells were kindly provided by Bob van de Water, Leiden University, The Netherlands. HCC1143 cells were cultured in RPMI 1640 Medium (ATCC modification) (Gibco) with 10% FBS (Sigma Aldrich). All cell lines tested negative for mycoplasma contamination. Cells were maintained at 37°C and 5 % CO2 in their respective growth medium without antibiotics. For hydrogel cultures, penicillin/streptomycin (Gibco) and Fungizone® (Gibco) were added to the regular culture medium at the time of cell plating.

### Hydrogel preparation and culture

Hydrogels used in this study were either prepared in-house (400Pa or 5000Pa) or purchased from Matrigen (500Pa or 8000Pa). In-house polyacrylamide-based hydrogels were prepared as described in Johnson, Leight, and Weaver[57] on 50 mm glass coverslips (VWR) or in 12-24 well glass bottom plates (MatTek). Briefly, glass surfaces were activated with 3-aminopropyltrimethoxysilane (Sigma Aldrich) followed by 0.5% glutaraldehyde (Sigma Aldrich). Solutions of 40% acrylamide and 2% bis-acrylamide (Both from BioRad) were mixed with 0.5M HEPES, pH 4.22, and H_2_O and *N*-succinimidyl ester of acrylamidohexanoic acid (N6 crosslinker, custom synthesized by Anthem Biosciences Pvt. Ltd., Bangalore India) was added. The final concentration of acrylamide/bis-acrylamide was 5.5%/0.15% for 5000 Pa gels and 3%/0.05% for 400 Pa gels. Solutions were degassed in a vacuum chamber (5305-0609 Nalgene) for 30-60 min before polymerization was initiated with N, N, N’, N’-tetramethylethylenediamine (TEMED, BioRad), and ammonium persulfate (Sigma Aldrich). A drop of polyacrylamide solution was put on the activated glass surface and a ClearVue rain clear (Turtle Wax) treated coverslip was carefully placed on top. Following polymerization, the top coverslip was removed, and the gel was conjugated with growth factor-reduced reconstituted basement membrane (rBM) (Corning or Biotechne, 0.14 mg/ml) in 50 mM HEPES, pH 8.0 for 4 hours on ice. The unreacted N6 crosslinker was then quenched in ethanolamine (1:100 in 50 mM HEPES, pH 8.0) for 30 min before gels were washed thoroughly in cold PBS. A final wash in cell culture media with penicillin/streptomycin and Fungizone® was then performed just before cell plating. Commercial gels (Matrigen, 0.5 kPa or 8 kPa) were conjugated with growth factor reduced rBM (Corning or Biotechne) (0.14 mg/ml in 50 mM HEPES, pH 8.0) o/n at 4°C before quenched, washed, and used as described for in-house gels. Media was changed to fresh culture media with pen/strep, fungizone, and rBM (0.2 mg/ml) every second day until the end of the experiment.

### Immunocytochemistry and confocal fluorescence imaging

Cells on hydrogels were fixed in 4% PFA (Sigma Aldrich) for 30 min at room temperature (RT) before permeabilizing in 0.2% Triton X-100 (Sigma Aldrich) in PBS for 15 min at RT. Cells were then blocked in 5% BSA in PBS for at least 1 h before incubated with primary antibodies overnight at 4°C. The following primary antibodies and dilutions were used; laminin (Serotec Ltd AHP420, diluted 1:200), integrin β4 (clone 3E1, Millipore MAB1964, diluted 1:200). Cells were washed twice in PBS and then incubated with appropriate secondary antibodies; AF647-conjugated anti-rabbit (Invitrogen A21245, 1:400) and/or AF488-conjugated anti-mouse (Thermo Scientific A-11029, 1:400) for 2 h at RT. Cells were counterstained with Hoechst 33342 (Sigma Aldrich 14533, 1:3000) and AF568-phalloidin (Thermo Scientific A-11029, 1:800). Images were acquired on a Nikon Ti2 inverted microscope with a Nikon 60x/1.2 water objective or a 20x/0.75 air objective. The microscope was equipped with a CrestOptics spinning disc (50um pinholes, 250um spacing), a back-illuminated sCMOS camera (Photometrics Prime 95B camera, 95% QE, 11um pixels), and the NIS-Elements software. The light source was a Lumencore Celesta and excitation and emission were as follows; 638nm excitation and 685/40nm emission to detect AF647-conjugated anti-rabbit; 477nm excitation and 511/20nm emission for AF488-conjugated anti-mouse; and 405nm excitation and 438/24 to detect Hoechst.

### PamGene peptide chip array

Commercially available peptide chip arrays from PamGene were used. All steps were performed according to the manufacturer’s protocol. Biological triplicates of CA1a cells were cultured on low (500Pa) or high (8 kPa) rBM-conjugated hydrogels (Matrigen) for three days. For integrin beta 1 inhibition, a monoclonal antibody (AIIB2, 5 µg/ml) was added to cells on high stiffness during the last 20 h of culture. Cells were washed twice in ice-cold PBS and lysed in M-PER mammalian extraction buffer (Pierce) containing Halt phosphatase and Halt protease inhibitor cocktails (Pierce) for 30 min on ice. Cell extracts were transferred to Eppendorf tubes and cleared by centrifugation at 13000 rpm for 15 min at 4°C. Protein concentrations were determined using BCA assay (Pierce) and aliquots were snap-frozen in liquid nitrogen and stored at -80°C. Measurements were performed on PamStation12 (PamGene) using 6 µg of protein extract for the PTK array and 2 µg for the STK array. The PTK array was processed in a single-step reaction where protein extract, ATP, and FITC-labelled pY20 antibody were incubated on the chip and the fluorescent detection of individual peptide phosphorylation was followed in real-time. The STK array was processed in two steps where the first step involved a 110-minute incubation with cell extracts, ATP, and primary antibody. In the second step, the secondary FITC-conjugated antibody was added, and the fluorescence signal was detected. Data were analyzed in the proprietary BioNavigator software (PamGene). Briefly, image analysis was performed to quantify fluorescent signals from each spot. A quality control step removed low signal spots and log2 transformed signals for remaining peptides were batch normalized using Combat correction[58, 59]. Differential peptide phosphorylation was then calculated as Log2 fold changes (high stiffness vs low and AIIB2 vs high) and statistically tested using an unpaired t-test. The list of differentially expressed peptides was finally used to predict upstream kinase activity via the Upstream Kinase Analysis tool, which ranks kinases based on a combined sensitivity and specificity score. Both scores are derived from permutation tests that assess whether the observed kinase activity change is statistically discernable and specific to the kinase’s target peptide set. Further detail on the PamGene peptide array based kinase prediction analysis can be found here: https://pamgene.com/wp-content/uploads/2024/08/Flyer_PamGene-Upstream-Kinase-Analysis-Tool.pdf.

### siRNA screen and image analysis

The siRNA screen was performed using in-house 5 kPa hydrogels in a 24-well plate format conjugated with rBM (0.14 mg/ml, Biotechne). MCF10CA1a cells were transfected with 20 nM SMART pool siRNA targeting the kinase of interest (Darmacon) using RNAiMAX (Invitrogen) and the manufacturer’s fast-forward protocol. A single FAK siRNA (target sequence 5’ GAA GTT GGG TTG TCT AGA A 3’) with confirmed knockdown efficiency was used as positive control and Allstars non-targeting control siRNA (Qiagen) was used as a negative control. 20 000 of transfected cells were plated on rBM-conjugated 5 kPa hydrogels. Overlay media containing 0.2 mg/ml rBM was added 24 h after transfection to allow 3D morphogenesis and cells were left in culture for three days before phenotype assessment. Cells were fixed in 4% PFA, permeabilized with 0.2% Triton X-100 and stained with AF568-phalloidin (1:800) and Hoechst (1:3000) in one single step. Large montage images with Z-stacks (5 planes, 32 µm total depth), covering the whole hydrogel area, were acquired using a Nikon Ti inverted microscope with a Nikon 10x/0.45 objective. The microscope was equipped with an Andor camera (Zyla 4.2+. fps at 2048×2048 pixels with pixel size 6.45 µm and QE 82%), a SpectraX LED light source, and the NIS-Elements software. Image analysis was performed with a custom GA3 pipeline in NIS-Elements to extract quantitative data. Segmentation was performed by thresholding the maximum intensity projections of red channel (actin), followed by quantification of the segmented area fraction of cell clusters. The area fraction was calculated as the ratio of the total area occupied by thresholded objects to the total imaged area. Measurements for the gene of interest (GOI) were normalized to the negative control (MOCK) on the same hydrogel plate (GOI/MOCK) and visualized using GraphPad Prism 10. Kinases whose knockdown resulted in a smaller area fraction than the positive control (FAK siRNA) were identified as regulators of the stiffness-induced malignant phenotype.

### Immunoblot

Cells were lysed using Strong RIPA buffer (10mM Tris-HCl-pH 7.5, 150mM NaCl, 1% Triton X100, 0.5% Na Deoxycolate, 0.1% SDS, 2mM EDTA) supplemented with protease (Roche) and phosphatase inhibitors (Sigma). The Lysates were clarified by centrifugation at 12,000 × g for 15 min at 4°C. Protein concentration was determined using the BCA assay (Pierce, Thermo Fisher). Equal amounts of protein (10–20 µg) were resolved by SDS-PAGE on 10% polyacrylamide gels (BioRad) and transferred onto PVDF membranes (Millipore) using a semi-dry transfer system (Bio-Rad). Membranes were blocked with 5% non-fat dry milk (PanReac AppliChem) in TBST (20 mM Tris, 150 mM NaCl, 0.1% Tween-20, pH 7.4) for 1 h at RT, followed by overnight incubation at 4°C with primary antibodies against IKBKE (Cell Signaling Technology, Cat. #D20G4, 1:1000), phosphorylated-MAPK8 (Novus Biologicals; Cat. #NB100-82009; 1:2000), or total-MAPK8 (Santa Cruz; Cat. #sc-1648; 1:5000). Membranes were then incubated with HRP-conjugated secondary antibodies (Jackson ImmunoResearch, 1:3000) for 1 h at RT. Bands were visualized using chemiluminescence (ECL) reagent (Pierce Thermo scientific) and imaged using an iBright Imaging System (Thermo). Quantification was performed using iBright Analysis Software. Local background corrected band volumes were quantified by the software and the values were normalized to loading controls, such as vincullin or total protein loading as visualized by Ponceau S (Sigma Aldrich) staining of the membranes to account for variations in sample loading.

### IKBKE and MAPK8 siRNA experiments

These experiments were performed using in-house prepared hydrogels (5 kPa) conjugated with 0.14 mg/ml Cultrex (Biotechne). The siRNAs were synthesized by Genepharma and the target sequences were as follows: IKBKE si1: 5’-TATCAAGCGTCCTTAGTCA -3’, IKBKE si2: 5’-GTACCTGCATCCCGACATG-3’, IKBKE si3: 5’-GCATTGGAGTGACCTTGTA-3’, IKBKE si4: 5’-GAACATCATGCGCCTCGTA -3’; MAPK8 si1: 5’-GCCCAGTAATATAGTAGTA-3’, MAPK8 si2: 5’-GGCATGGGCTACAAGGAAA-3’, MAPK8 si3: 5’-GAATAGTATGCGCAGCTTA-3’, and MAPK8 si4: 5’-GATGACGCCTTATGTAGTG-3’. In these experiments, transfection of 20 nM of siRNA was performed at the time of plating cells on hydrogels using RNAiMAX (Invitrogen) according to the manufacturer’s fast-forward protocol. AllStars Negative Control siRNA (Qiagen) was used as a negative control. Media was changed 16-20h after transfection and again 6 h before fixation. Phenotype and Edu positive cell fractions were assessed 4 d after siRNA transfection.

### IKBKE and MAPK8 inhibitor experiments

These experiments were performed using in-house prepared 5 kPa hydrogels conjugated with 0.14 mg/ml Cultrex (Biotechne). The IKBKE inhibitor Amlexanox (Abcam) and MAPK8 inhibitor JNK-IN-8 (Sigma Aldrich), used at concentrations and time indicated in the figure legends. Phenotype and Edu positive cell fractions were compared to DMSO (vehicle) treated cells.

### Cell proliferation assay

Cells were cultured on in-house prepared 5 kPa hydrogels for the specified time, then incubated at 37°C for 1 h with 40µM Edu (Invitrogen). After fixation, EdU incorporation was detected using a Click-iT Plus EdU Alexa Fluor 488 Flow Cytometry Assay Kit (Invitrogen) following the manufacturer’s protocol. Images were acquired using a Nikon Ti inverted microscope with a Nikon 20x/0.75 objective. The microscope was equipped with an Andor Zyla 4.2+ camera (fps at 2048x2048 pixels with pixel size 6.45 µm and QE 82%), a SpectraX LED light source, and the NIS-Elements software. The fraction of EdU positive cells relative to total cells was analyzed using CellProfiler 4.2.6.

### Statistical analysis

All data visualization and statistical analysis were performed in GraphPad Prism software version 10 and the statistical tests are stated in figure legends.

## Acknowledgements

Imaging was performed at the Live Cell Imaging Core Facility and Nikon Center of Excellence at Karolinska Institutet. This study was supported by grants to SS from The Swedish Research Council (2021-01139); The Swedish Cancer Society (23 2856 Pj); and Radiumhemmets forskningsfonder (201353). Anagha Acharekar was supported by a Scholarship from The Wenner-Gren Foundation.

## Author contributions

F.Y. conceived, designed, and analyzed the siRNA screen; conceived, designed, partially conducted, and analyzed IKBKE and MAPK8 experiments; and wrote the manuscript. S.G. conceived, designed, and conducted model characterization experiments. S.G. conceived, designed, conducted, and analyzed the peptide array experiments with help from PamGene, The Netherlands; and wrote the manuscript.

H.O. performed model characterization experiments, siRNA screen experiments, and IKBKE-related experiments. C.V. and A.A. performed immunoblot experiments. S.S. conceived and supervised the study, provided funding, and edited the manuscript. All authors approved the final manuscript.

## Figure legends

**Supplementary Fig. 1.**
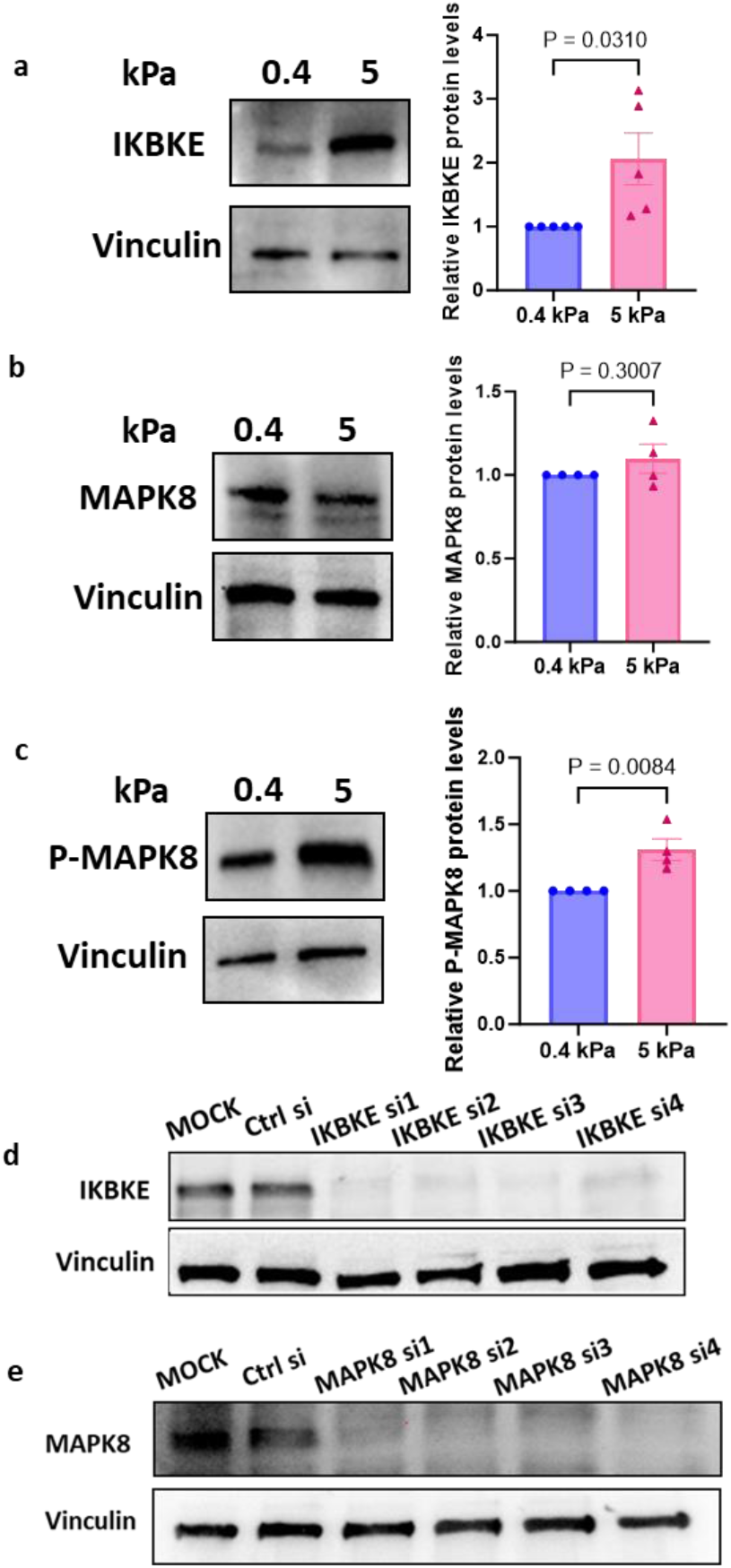

**Supplementary Fig. 2.**
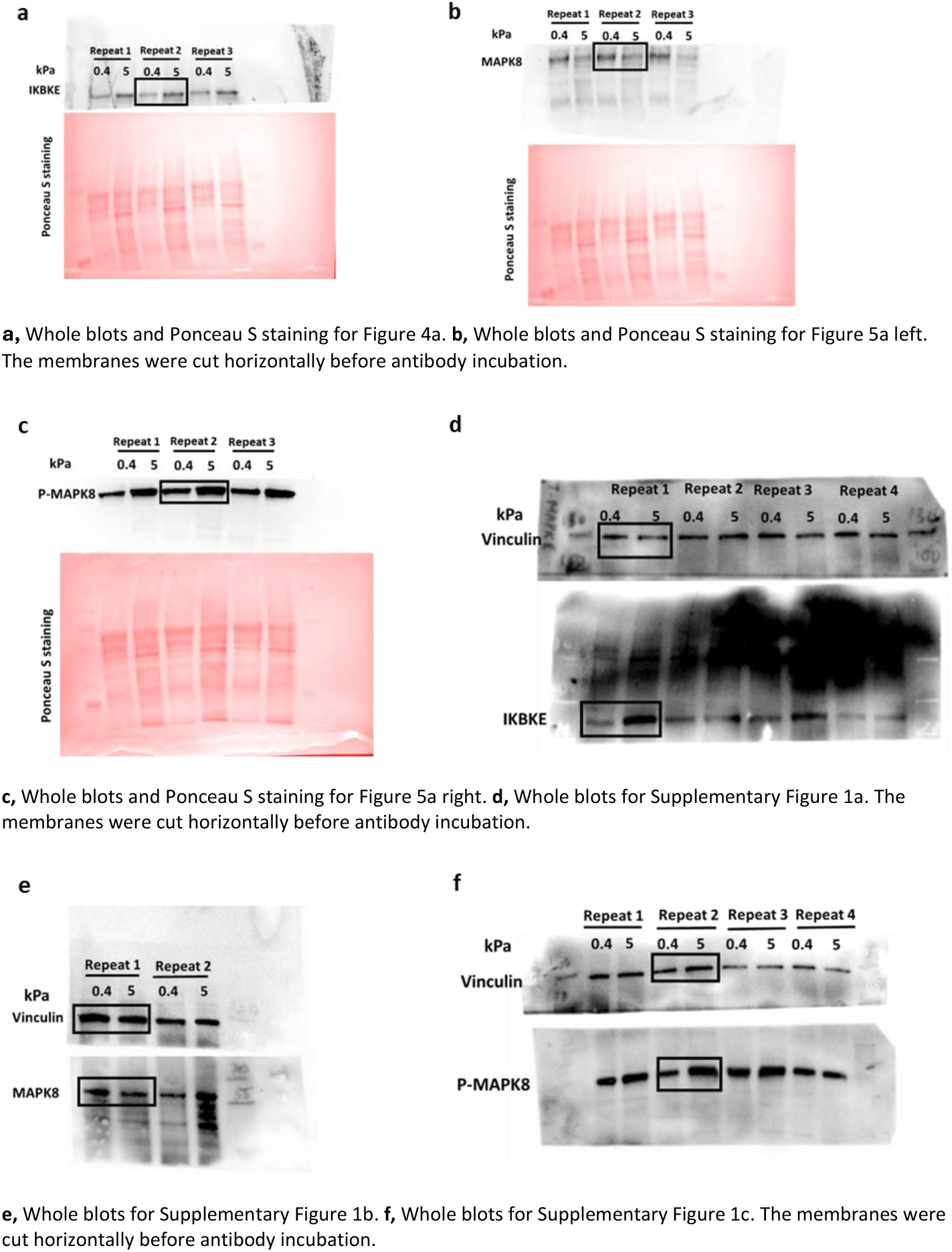
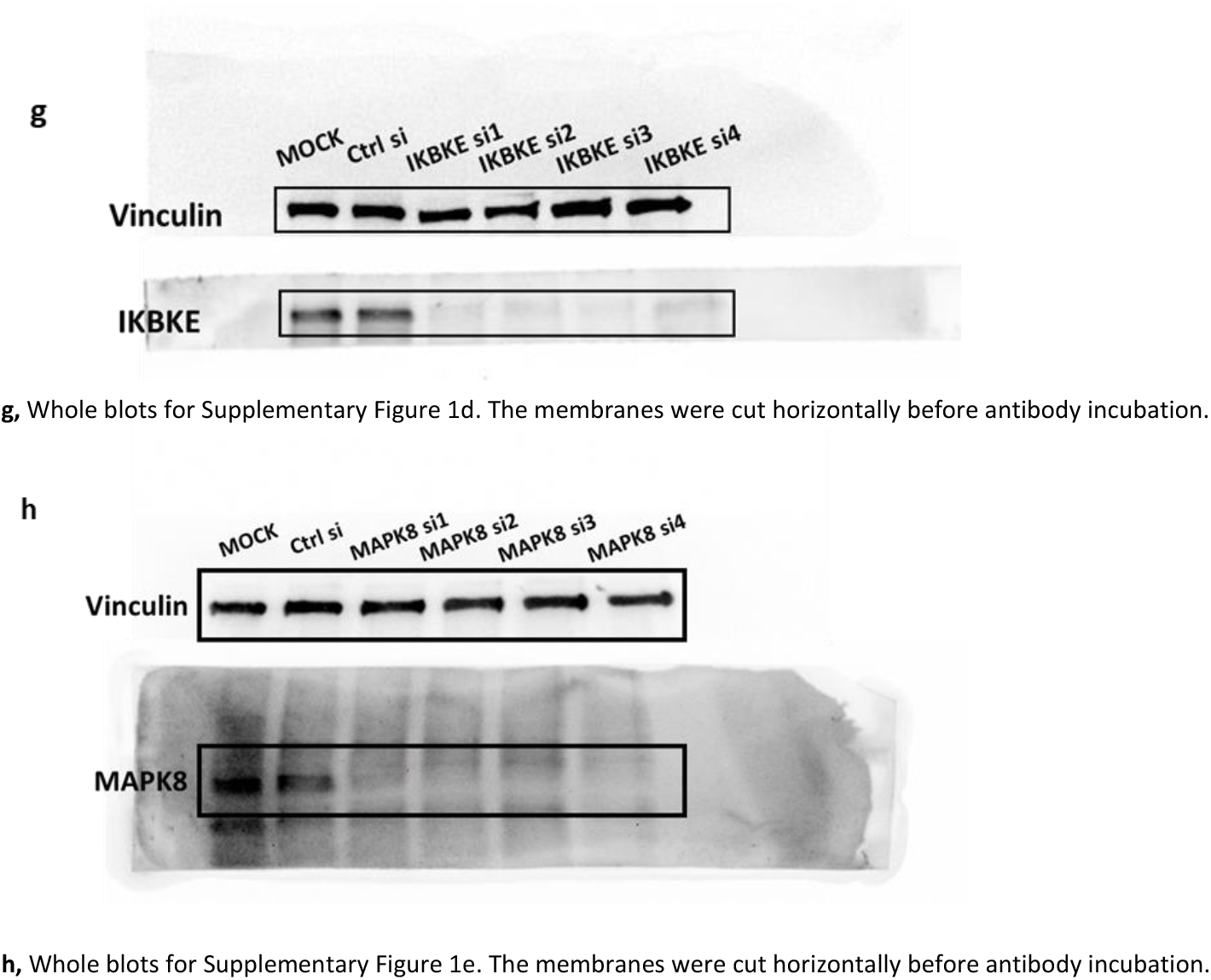

